# Resistance to EGFR inhibitors in lung cancer occurs through horizontal transfer and is associated with increased caveolins expression

**DOI:** 10.1101/2022.02.20.481179

**Authors:** Susana Junqueira-Neto, Ana R. Oliveira, Joana F. Marques, Patrícia Oliveira, Maria G. Fernandes, Venceslau Hespanhol, Ana Barroso, Jorge Pinheiro, Conceição Souto-Moura, Bruno Cavadas, Luísa Pereira, Bárbara Adem, Miguel Silva, Sónia A. Melo, José L. Costa, José C. Machado

## Abstract

Resistance to treatment is a major clinical problem and a major cause of cancer-related deaths. Understanding the biological basis of resistance acquisition is of utmost importance to improve the clinical management of cancer patients. NGS analysis of human lung cancer (LC) tumors from patients that relapsed after treatment with EGFR-tyrosine kinase inhibitors (TKI), revealed that the p.T790M resistance mutation is not present in all the relapsing tumor cells, suggesting that LC cells can become resistant even if not carrying the p.T790M mutation. Using in vitro treatments with conditioned medium (CM) and in vivo co-inoculation experiments, we show that LC cells sensitive to EGFR-TKIs (S cells) acquire resistance faster when treated with CM from LC cells resistant to EGFR-TKIs (R cells) or when co-inoculated with R cells in opposite flanks of the same animal. Importantly, we show that acquisition of resistance is not due to the emergence of subpopulations of cancer cells with new resistance mutations. Using transcriptomics, we show that acquisition of resistance is associated with upregulation of genes involved in endocytosis, namely caveolins CAV1 and CAV2. These findings were validated in human clinical samples, where an increase in CAV1 and CAV2 expression was associated with tumor relapse after treatment with EGFR-TKIs. Our results suggest that acquisition of resistance to targeted therapies results from the combined effect of selection of cells harboring specific resistance mutations and horizontal transfer of the resistance phenotype. These findings may pave the way to bring intercellular communication into the realm of cancer treatment.

**One Sentence Summary:** Resistance to EGFR inhibitors is transferred horizontally between lung cancer cells and is associated with gain of expression of caveolins.

## INTRODUCTION

Targeted cancer therapies block specific signalling pathways implicated in proliferation and survival of cancer cells *(1, 2)*. Despite increasing progression-free and overall survival of cancer patients, resistance to these drugs is almost universally observed *(3)*. Thus, acquired resistance to therapy is a major clinical problem and a cause of therapy failure *(4)*. Acquisition of resistance to targeted therapies is currently explained by the selective accumulation of cancer cells carrying resistance-conferring mutations *(5, 6)*. However, data on record sustain that cell division and selection cannot by itself explain the speed at which a tumour becomes clinically resistant *(7, 8)*. A study about disease progression of a melanoma patient with the BRAF p.V600E mutation is a paradigmatic example *(9)*. Furthermore, clinical data suggests that in lung cancer (LC) patients who relapse after EGFR tyrosine kinase inhibitor (TKI) treatment, cells that carry and cells that do not carry a specific resistance-conferring mutation may coexist in a treatment-resistant tumour *(10, 11)*. Hence, acquisition of resistance to targeted therapies may be influenced by mechanisms of horizontal transfer, complementing a vertical transfer model entirely dependent on cell division and selection of cells carrying resistance mutations.

Lung cancer is a prototypical model of successful use of targeted therapies, where EGFR kinase domain mutations hyperactivate and confer dependency on the EGFR oncogenic pathway for cell survival *(12, 13)*. Treatment of EGFR-mutant LC with EGFR-TKIs leads to successful clinical response in many patients *(14, 15)*. Despite initial benefit, disease progression typically develops after 9-12 months of treatment, because of the recurrent EGFR mutation p.T790M in 50-60% of cases *(16-18)*. The p.T790M mutation increases the affinity of EGFR to adenosine triphosphate, relative to its affinity to TKIs *(19)*. Resistance to EGFR-TKI therapy may also be achieved through mutations in other genes, such as MET and ERBB2 *(20, 21)*, showing that therapy resistance does not depend solely on interference with drug activity, but may elapse also from alternative cell signalling cues. Therefore, LC constitutes a prime model to address the issue of resistance acquisition to targeted therapies, given that both sensitivity and resistance-conferring mechanisms have been identified.

We show that LC cells that do not carry known EGFR-TKI resistance mutations, acquire resistance to erlotinib faster when in the presence of LC cells that carry the p.T790M EGFR mutation. We also show that acquisition of resistance is associated with increased expression of the endocytosis associated proteins CAV1 and CAV2. Our study provides novel insight on how therapy resistance becomes a predominant phenotype in cancer and shows that widespread expression of a therapy resistance phenotype is not strictly dependent on division and selection of subpopulations of cancer cells carrying resistance triggering mutations.

## RESULTS

### Human lung cancers present the p.T790M resistance mutation only in a fraction of the relapsing tumour cells

Studies that characterize the EGFR sensitizing and resistance mutations *(10, 11)*, show that in some patients the allele frequency (AF) of the initial sensitizing mutation is higher than that of the resistance-conferring mutation. This suggests the resistance mutation is present only in a fraction of the relapsing tumour cells. Because this had never been studied in a systematic way, we used an NGS panel to analyse a series of 28 samples, including 14 liquid and 14 tissue biopsies (table 1), from LC patients who relapsed after EGFR-TKI treatment and where the p.T790M resistance mutation was detected.

**Table 1.** Allele frequencies determined by NGS in LC samples after disease progression.

| Patient | Biopsy type | EGFR sensitizing mutation | AF sensitizing mutation | AF p.T790M mutation | Fraction |
| --- | --- | --- | --- | --- | --- |
| 1 | Liquid | c.2239_2256del; p.Leu747_Ser752del | 0.8% | 0.3% | 62.5% |
| 2 | Liquid | c.2573T>G; p.Leu858Arg | 65% | 29% | 55.4% |
| 3 | Liquid | c.2235_2249del; p.Glu746_Ala750del | 13% | 3% | 76.9% |
| 4 | Liquid | c.2235_2249del; p.Glu746_Ala750del | 3% | 0.3% | 90.0% |
| 5 | Liquid | c.2573T>G; p.Leu858Arg | 19% | 3% | 84.2% |
| 6 | Liquid | c.2573T>G; p.Leu858Arg | 11% | 7% | 36.4% |
| 7 | Liquid | c.2235_2249del; p.Glu746_Ala750del | 3% | 1.8% | 40.0% |
| 8 | Liquid | c.2236_2250del; p.Glu746_Ala750del | 2.6% | 0.8% | 69.2% |
| 9 | Liquid | c.2236_2250del; p.Glu746_Ala750del | 11.9% | 1.7% | 85.7% |
| 10 | Liquid | c.2573T>G; p.Leu858Arg | 27.6% | 1% | 96.4% |
| 11 | Liquid | c.2240_2257del; p.Leu747_Pro753delInsSer | 2% | 0.6% | 70.0% |
| 12 | Liquid | c.2235_2249del; p.Glu746_Ala750del | 11% | 4.2% | 61.8% |
| 13 | Liquid | c.2240_2257del; p.Leu747_Pro753delInsSer | 1.8% | 1.9% | 0% |
| 14 | Liquid | c.2235_2249del; p.Glu746_Ala750del | 82% | 47% | 42.7% |
| 15 | Tissue | c.2238_2261delInsGCAAC; p.Leu747Glnfs*13 | 61% | 10% | 83.6% |
| 16 | Tissue | c.2573T>G; p.Leu858Arg | 20% | 6% | 70.0% |
| 17 | Tissue | c.2240_2257del; p.Leu747_Pro753delInsSer | 24% | 18% | 25.0% |
| 18 | Tissue | c.2235_2249del; p.Glu746_Ala750del | 29% | 7% | 75.9% |
| 19 | Tissue | c.2235_2249del; p.Glu746_Ala750del | 40% | 14% | 65.0% |
| 20 | Tissue | c.2235_2249del; p.Glu746_Ala750del | 84% | 10% | 88.1% |
| 21 | Tissue | c.2573T>G; p.Leu858Arg | 53% | 25% | 52.8% |
| 22 | Tissue | c.2235_2249del; p.Glu746_Ala750del | 31% | 8% | 74.2% |
| 23 | Tissue | c.2573T>G; p.Leu858Arg | 26% | 2% | 92.3% |
| 24 | Tissue | c.2573T>G; p.Leu858Arg | 41% | 1% | 97.6% |
| 25 | Tissue | c.2235_2249del; p.Glu746_Ala750del | 45% | 24% | 46.7% |
| 26 | Tissue | c.2240_2257del; p.Leu747_Pro753delInsSer | 15% | 3% | 80.0% |
| 27 | Tissue | c.2236_2250del; p.Glu746_Ala750del | 60% | 16% | 73.3% |
| 28 | Tissue | c.2236_2250del; p.Glu746_Ala750del | 14% | 7% | 50.0% |

AF, allele frequency; Fraction, the fraction of cancer cells that do not carry the resistance mutation as inferred by the formula $(1 - \text{AF p.T790M} / \text{AF sensitizing})$ .

The AF of the initial sensitizing EGFR mutation was always higher than that of the p.T790M mutation in the relapsing tumours, except for one case (table 1 and Fig. S1, P<0.0001). On average, the AF of the initial mutation was 3.2-fold higher than that of the p.T790M mutation, irrespective of the sample being a liquid or tissue biopsy and irrespective of the type of initial EGFR mutation, namely being a point mutation or an indel, which could affect read counting and comparison using NGS. No additional EGFR-TKI resistance mutations in any of the genes analysed were detected in these cases, excluding co-occurrence of resistance mutations as the cause for p.T790M-independent resistance. Likewise, no CNVs in EGFR were detected, excluding ploidy as the cause for observed differences in AF.

The fact that in human LC we frequently observe a lower AF for the p.T790M mutation in comparison with the sensitizing mutation in the same tumour, suggests that a fraction of the cancer cells “borrow” their resistance phenotype from cells that carry the p.T790M mutation. Our results suggest that the fraction of cancer cells that do not carry the resistance mutation may be higher than 90% in some tumors (table 1).

### Resistant cells accelerate *in vitro* acquisition of resistance by sensitive cells to EGFR-TKI

Since cancer cells can display an EGFR-TKI resistant phenotype without necessarily carrying a resistance mutation, we assessed whether resistant cells could influence acquisition of resistance to EGFR-TKI by sensitive cells *in vitro*. We used the erlotinib-sensitive LC cell line HCC827 (S cells) that harbours the sensitizing EGFR p.E746-A750del mutation, and the erlotinib-resistant LC cell line H1975 (R cells) that harbours both a sensitizing and a resistance mutation (EGFR p.L858R and p.T790M, respectively).

S cells treated with CM from R cells (Fig. 1A) achieved resistance faster in comparison to non-treated S cells (90±3 vs. 120±5 days, P=0.0009, Fig. 1B). Follow-up of S cells that acquired resistance and were further kept under erlotinib treatment only, shows that once the resistance phenotype is acquired, cells maintained a stable resistance phenotype until we stopped follow-up at 55 days.

**Figure 1.**
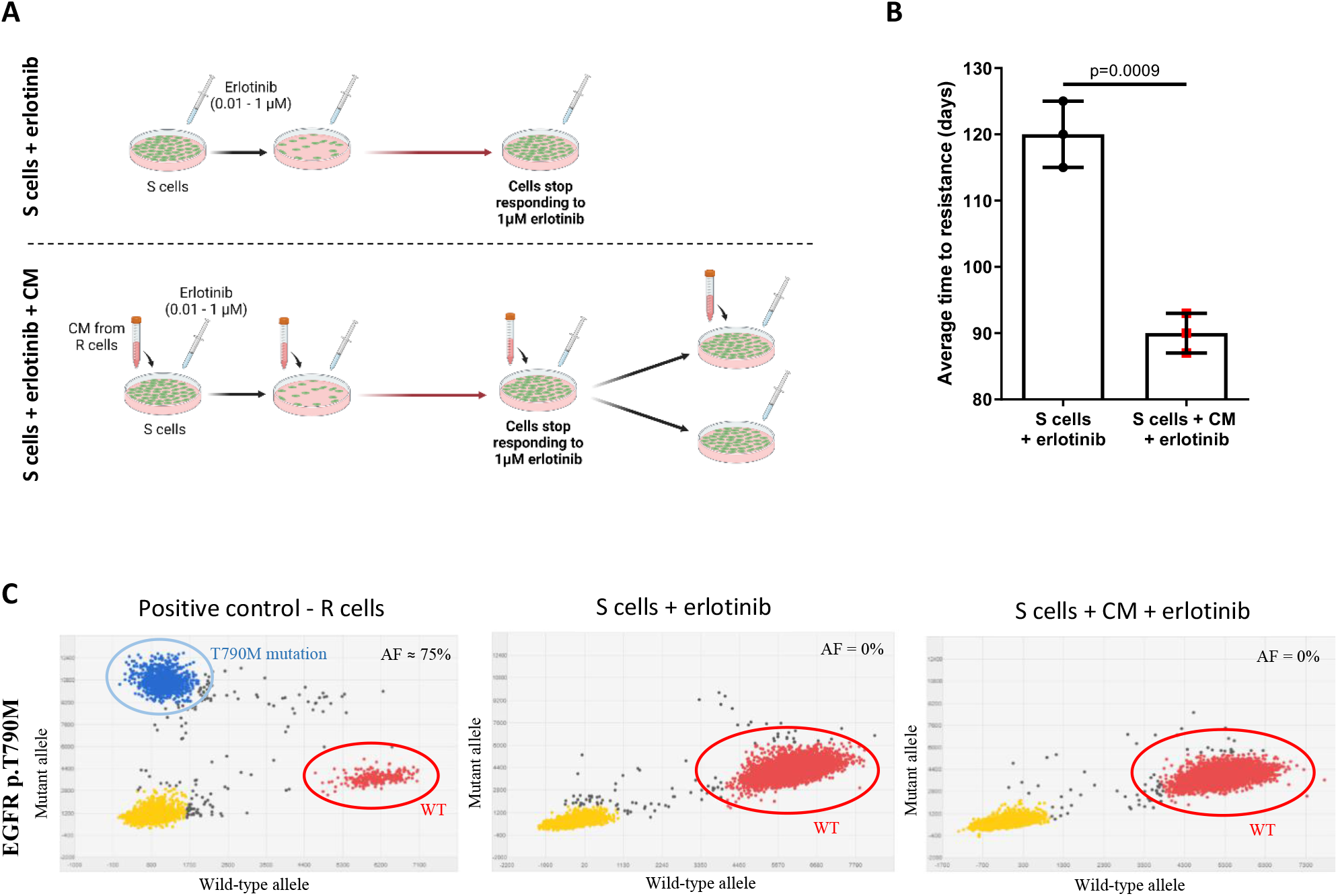
R cells accelerate *in vitro* acquisition of resistance to EGFR-TKI by S cells. (**A**) Experimental outline of S cells treated with conditioned media (CM) from R cells, and S cells without CM. In both experiments, S cells were treated with increasing concentrations of erlotinib (0.01-1 μM). Resistance was achieved when cells stopped responding to the treatment. CM-treated cells were further divided into two groups, one maintaining the treatment with CM and erlotinib, and the other with erlotinib only. (**B**) Comparison of the average time to resistance in days between CM-treated cells and CM-untreated cells (Mean ± SD). (**C**) Digital PCR plots for the p.T790M assay. The *EGFR* p.T790M mutation is not present in DNA from any of the CM-treated or CM-untreated cells: yellow dots represent empty wells; blue dots represent the T790M variant; red dots correspond to the wild-type sequence; and grey dots represent polyclonal wells not considered for analysis. The AF (allelic frequency) is calculated dividing the number of blue dots by the sum of blue and red dots.

Genetic analysis of DNA from S cells revealed that the p.T790M mutation, present in R cells, could not be detected in S cell cultures, confirming that no cell contamination occurred, and suggesting that acquisition of resistance to erlotinib in S cells is not strictly dependent on the presence of the p.T790M mutation (Fig. 1C).

### Resistant cells accelerate *in vivo* acquisition of resistance by sensitive cells to EGFR-TKI

To evaluate whether R cells could influence acquisition of resistance to EGFR-TKI by S cells *in vivo*, we used a strategy of single (SI) or dual inoculation (DI) of LC cells in Rag2-/-;IL2rg-/- immunodeficient mice (Fig. 2A). Our results show that S tumours in DI (S and R cells) mice acquire resistance significantly faster in comparison to SI (S cells only) mice (78±5 vs. 132±16 days, P=0.0006, Fig. 2B and 2C), which is reflected in a significant decrease in progression-free survival in the DI group (Fig. 2D, P=0.001). S tumours with acquired resistance to erlotinib were transplanted into new mice for two passages, always under erlotinib treatment, and maintained the ability to grow (Fig. 2E), reinforcing the *in vitro* observation that acquisition of resistance results in a stable phenotype.

**Figure 2.**
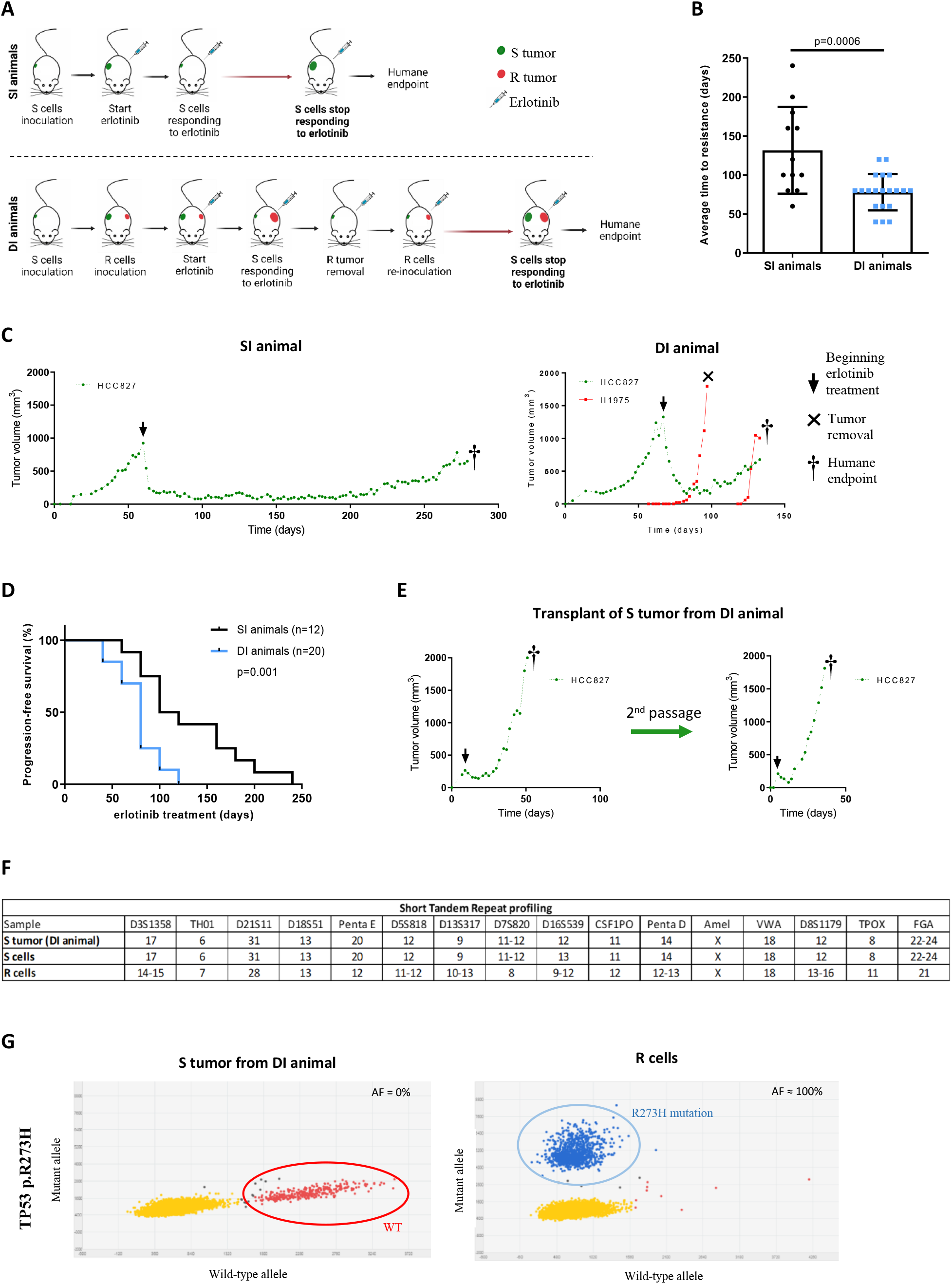
R cells accelerate *in vivo* acquisition of resistance to EGFR-TKI by S cells. (**A**) Experimental outline of mice engrafted with S cells inoculated in a single flank (SI animals) and mice engrafted with S and R cells inoculated in opposite flanks (DI animals) treated with erlotinib 3 times/week by oral gavage. (**B**) Comparison of the average time to relapse in days between DI and SI animals (Mean ± SD). (**C**) Tumor growth kinetics of one representative SI and one representative DI mouse, illustrating that S tumors in DI animals acquire resistance faster. (**D**) Progression-free survival curves of SI and DI animals are significantly different (*P*=0.001; Log-rank Mantel-Cox test). (**E**) After resistance acquisition, S tumors from DI animals were re-inoculated into new mice for two passages and erlotinib treatment was maintained during the whole experiment. (**F**) Comparison of the STR profile of S tumors from DI animals with the STR profile of S and R cells. (**G**) dPCR plots of S tumor from a DI animal negative for the *TP53* p.R273H mutation and of R cells positive for the *TP53* p.R273H mutation with 100% AF.

To discard the possibility that migrating R cells could explain the acquisition of resistance to erlotinib in S tumors, we performed genetic analysis. Short tandem repeat (STR) profiling of S and R cells, and of S tumours from DI animals showed a perfect overlap between the profile of S tumours and S cells (Fig. 2F). To maximize the sensitivity of detection of potential migrating R cells, we also performed digital PCR for the TP53 p.R273H mutation that is present in R cells with an AF of 100% and could not detect it in S tumours (Fig. 2G). Histology analysis of R and S tumours also shows distinct morphological patterns compatible with distinct cellular origin of the tumours (Fig. S2).

Using NGS, we identified the EGFR p.E746_A750del mutation in every S tumour from DI animals with an allelic frequency (AF) similar to that of the parental S cell line (≈90%; table S1). In contrast, the p.T790M mutation is detected only in 6 out of 20 tumours and always with very low AF (≈0.1%). These results were confirmed by dPCR (Fig. 3A) and corroborate the *in vitro* observation that acquisition of resistance to erlotinib in S tumors is not dependent on the presence of the p.T790M mutation. The vestigial AF of p.T790M is most likely the result of circulating cfDNA derived from R cells growing in the opposite flank of the mice since we could not detect the p.T790M mutation in any of the S tumours from SI animals (table S1). To verify this, we analyzed liquid biopsies from mice where R cells had been inoculated and confirmed that the p.T790M mutation can indeed be detected in circulating cfDNA (Fig. 3B).

**Figure 3.**
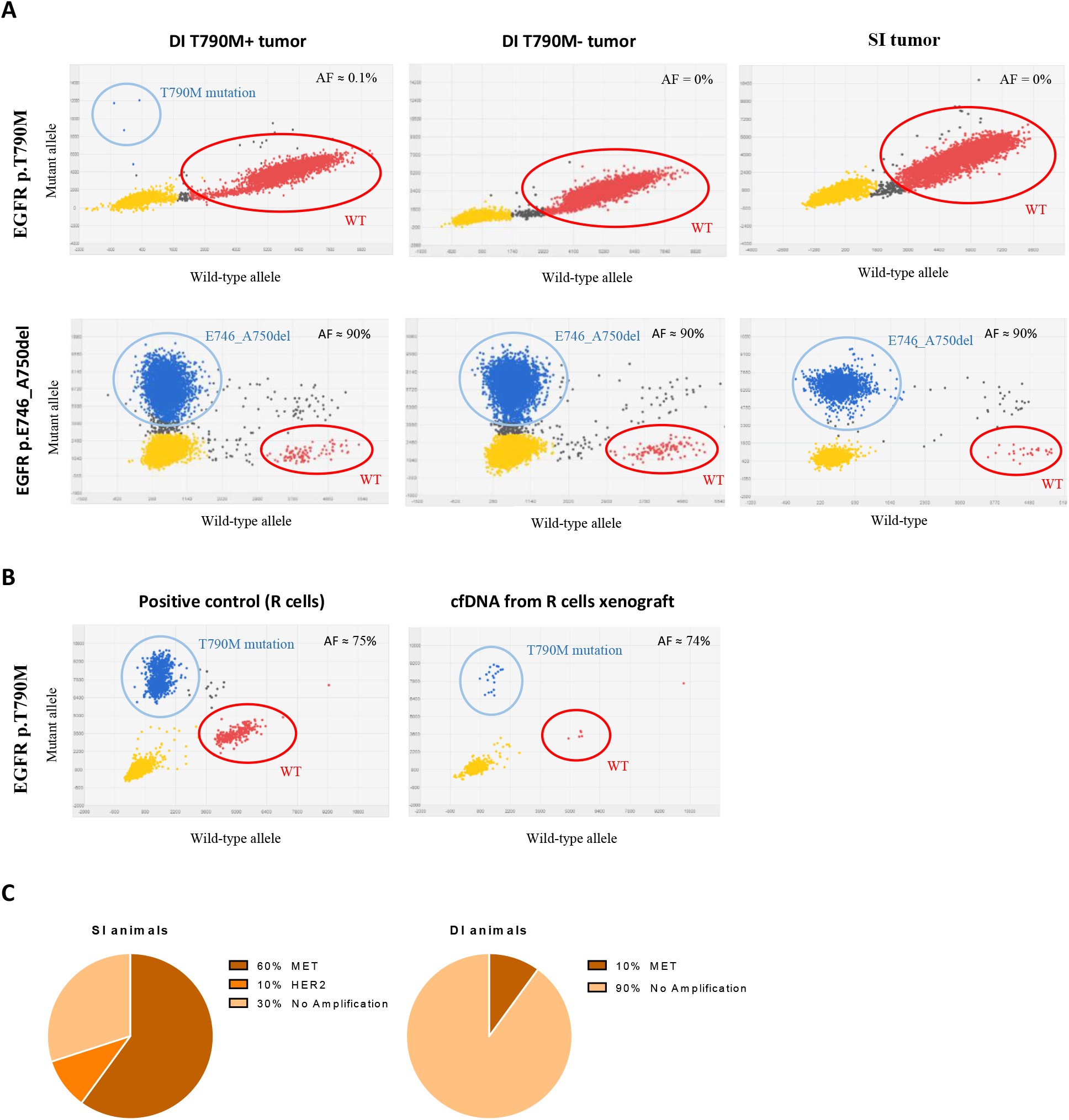
Detection of resistance mutations to EGFR-TKI in S tumors from SI and DI animals. (**A**) dPCR plots for *EGFR* p.T790M and *EGFR* p.E746_A750del mutations in representative examples of S tumors from SI and DI animals. S tumor from DI animal positive for *EGFR* p.T790M mutation with 0.1% AF (left panel); S tumor from DI animal negative for *EGFR* p.T790M mutation (middle panel); S tumor from SI animal negative for *EGFR* p.T790M mutation (right panel). All samples are positive for the *EGFR* p.E746_A750del mutation with ≈90% AF. (**B**) dPCR plots for the p.T790M mutation. The *EGFR* p.T790M mutation is present in the cfDNA from mice xenografted wit R cells (right panel) with an AF similar to that of the parental R cell line (left panel). (**C**) NGS data on *MET* and *HER2* gene amplification in S tumors from SI animals (left diagram) and from DI animals (right diagram).

To verify whether other mutations that confer resistance to EGFR-TKIs were present in resistant tumours, we used an NGS panel encompassing all known resistance mutations, including point mutations in EGFR, KRAS, NRAS, BRAF and PIK3CA, and CNVs in HER2, MET and BRAF. No additional point mutations were identified. Interestingly, in S tumours from SI animals we identified HER2 or MET amplification in 70% of the cases (6 and 1, respectively), indicating that mutation-driven resistance is a prevalent mechanism in these tumours (Fig. 3C). However, in S tumours from DI animals, only MET amplification was found and only in two cases (10%, Fig. 3C). These results suggest that mutation-driven resistance, through the EGFR p.T790M or any other known EGFR-TKI resistance mutation, is not prevalent in S tumours that acquire resistance to erlotinib in DI mice.

### Caveolins are overexpressed in tumours from DI animals that acquire resistance to EGFR-TKI

Since the presence of resistance mutations does not seem to be a pre-requisite for acquisition of resistance to erlotinib in S tumors from DI animals, we sought to identify what differentiates S tumours growing in the presence or absence of R tumours. We used RNA-Seq to identify differentially expressed genes in S tumours not treated with erlotinib and S tumors from DI animals treated with erlotinib. In other words, we compared erlotinib-resistant S tumors (ERS) to erlotinib-sensitive S tumors (ESS). Differentially expressed genes in ERS and ESS tumours were also compared with the expression profile of R tumours under the assumption that genes relevant to explain resistance to erlotinib in this specific biological context should be shared between ERS and R cells.

Transcriptome analysis showed that ERS tumours cluster together and are significantly different from ESS tumours (Fig. 4A). Out of the 2642 differentially expressed genes (FDR P<0.05), we represented the top 10 differentially expressed genes (Fig. 4B). Using a gene ontology approach, we observed that endocytosis was the KEGG pathway with the highest number of differentially expressed genes detected (Fig. 4C, 55 genes, P=2.46E-0.3). Within the endocytosis pathway, CAV1 and CAV2 were the most significantly overexpressed genes in ERS tumours in comparison with ESS tumours (CAV1: fold change=7.7x, FDR P=1.23E-0.5; and CAV2: fold change=4.2x, FDR P=4.9E-0.7, Fig. 4D and Fig. S3).

**Figure 4.**
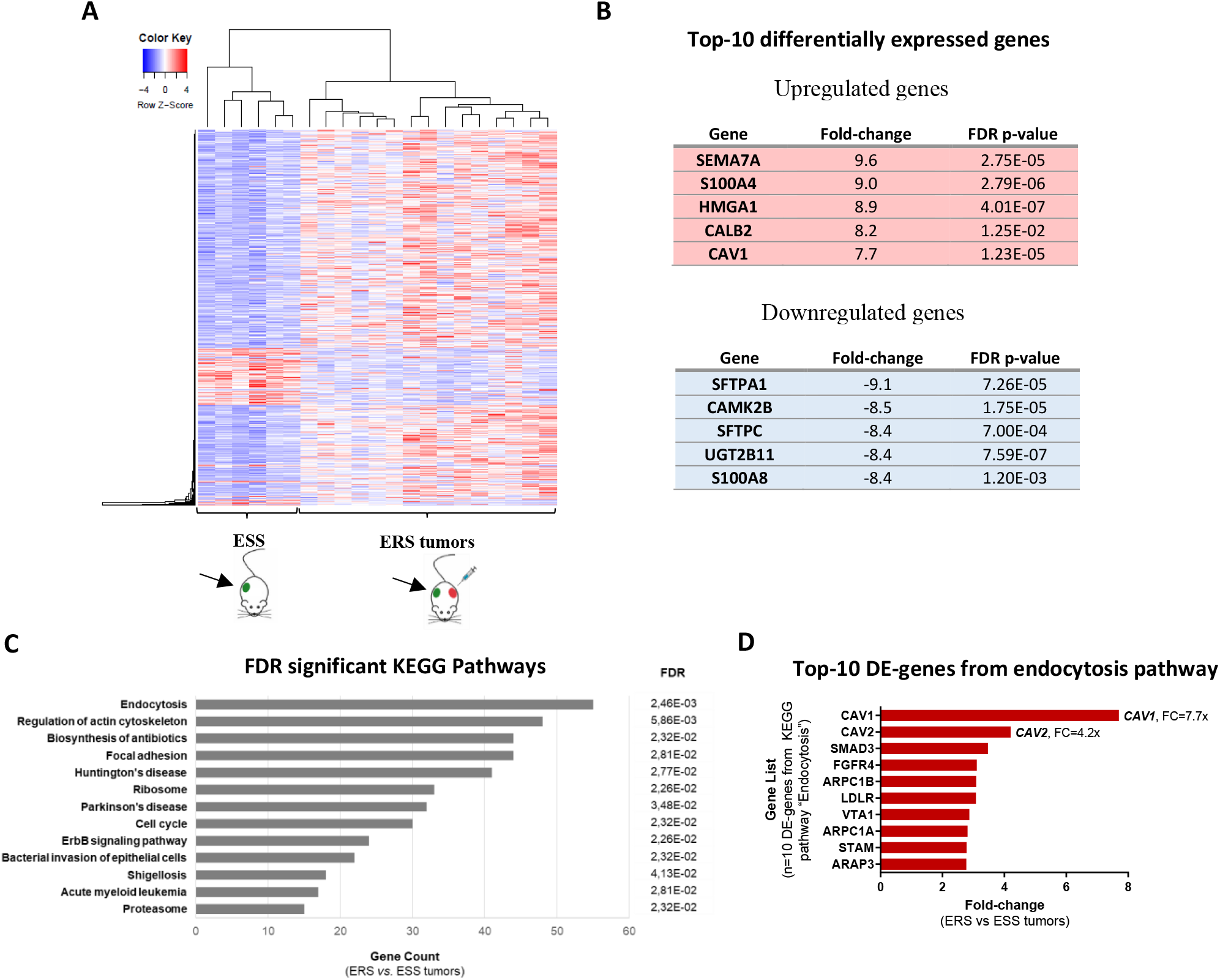
Identification of endocytosis as the mechanism associated with resistance acquisition in ERS tumors. (**A**) Heatmap for RNA-Seq gene expression profiles for ESS tumors versus ERS tumors. FDR p-value <0.05; 2642 differentially expressed (DE) genes. (**B**) Top-10 DE-genes between ERS and ESS tumors (top-5 upregulated genes and top-5 downregulated genes). (**C**) Gene ontology showed endocytosis KEGG pathway as the pathway with the highest number of genes detected and the best FDR p-value (55 genes, FDR *P*=2.46E-0.3). (**D**) Top-10 DE-genes between ERS and ESS tumors from the endocytosis KEGG pathway (FDR *P*<0.05). FC – fold change.

Expression analysis through qPCR validated that CAV1 and CAV2 are overexpressed both in ERS tumours and R tumours, when compared with ESS tumours (Fig. 5A). The same was observed for protein immunohistochemistry expression, with both ERS tumours and R tumours showing strong CAV1 and CAV2 expression in tumours cells, whereas in ESS tumours CAV1 and CAV2 expression showed weak and focal expression in tumours cells (Fig. 5B and C).

**Figure 5.**
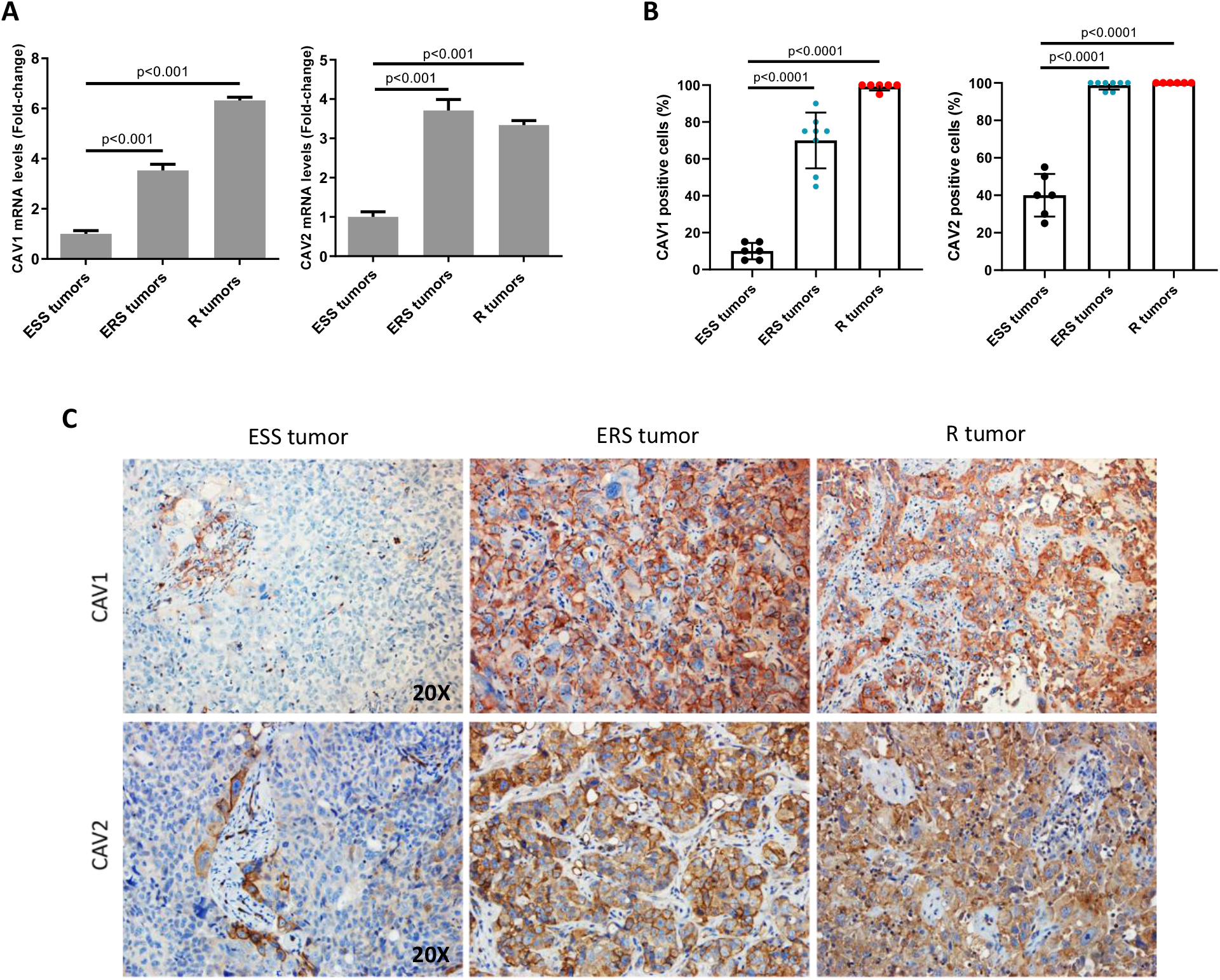
Validation of increased caveolins expression in ERS tumors by real time PCR and immunohistochemistry (IHC). (**A**) The bars represent the 2^-ΔΔCt^ as a fold-change of the *CAV1* (left) and *CAV2* (right) mRNA expression, normalized to *GUSβ* and *HPRT1*, for ESS tumors, ERS tumors and R tumors. (Mean ± SE; *P*<0.001). (**B**) Percentage of CAV1 and CAV2 protein positive cells for ESS tumors, ERS tumors and R tumors (Mean ± SD; *P*<0.0001). (**C**) CAV1 and CAV2 immunohistochemistry expression in representative examples of ESS tumors, ERS tumors and R tumors (magnification x200). ESS tumors show weak focal expression in tumor cells and staining in perivascular cells; ERS tumors and R tumors show strong and diffuse expression of caveolins in tumor cells.

### Caveolins are overexpressed in human lung cancers that progress after EGFR-TKI treatment

To validate our findings, we compared the immunohistochemical expression of CAV1 and CAV2 in human LC before and after treatment with erlotinib. We selected fifteen cases of LC harbouring sensitizing EGFR mutations and treated first-line with erlotinib, in which the p.T790M EGFR mutation was detected as the resistance mechanism, and for which treatment-naïve and post-progression tissue biopsies were available. CAV1 and CAV2 protein expression was mainly localized in the cell membrane of tumour cells, although some cytoplasmic staining could be detected as well. Staining in the endothelium and connective tissue was also detected (Fig. 6A). Out of the 15 pairs of samples analysed for CAV1 expression, 80% were negative before treatment and 20% showed moderate expression (Fig 6B). CAV2 expression could only be analysed in 10 pairs of cases due to lack of material. 75% of the cases were negative before treatment, 17% displayed moderate expression and 8% had strong expression (Fig 6B). From pre-treatment to post-progression samples, we observed a significant increase (P=0.007) in CAV1 expression in 67% of the cases: from no to moderate expression in 5 cases; from no to strong expression in 3 cases; and from moderate to strong expression in 2 cases (Fig. 6B, table S2). A significant increase (P=0.04) in CAV2 expression was observed in 60% of the cases: from no to moderate expression in 3 cases; from no to strong expression in 1 case; and from moderate to strong expression in 2 cases (Fig. 6B, table S2).

**Figure 6.**
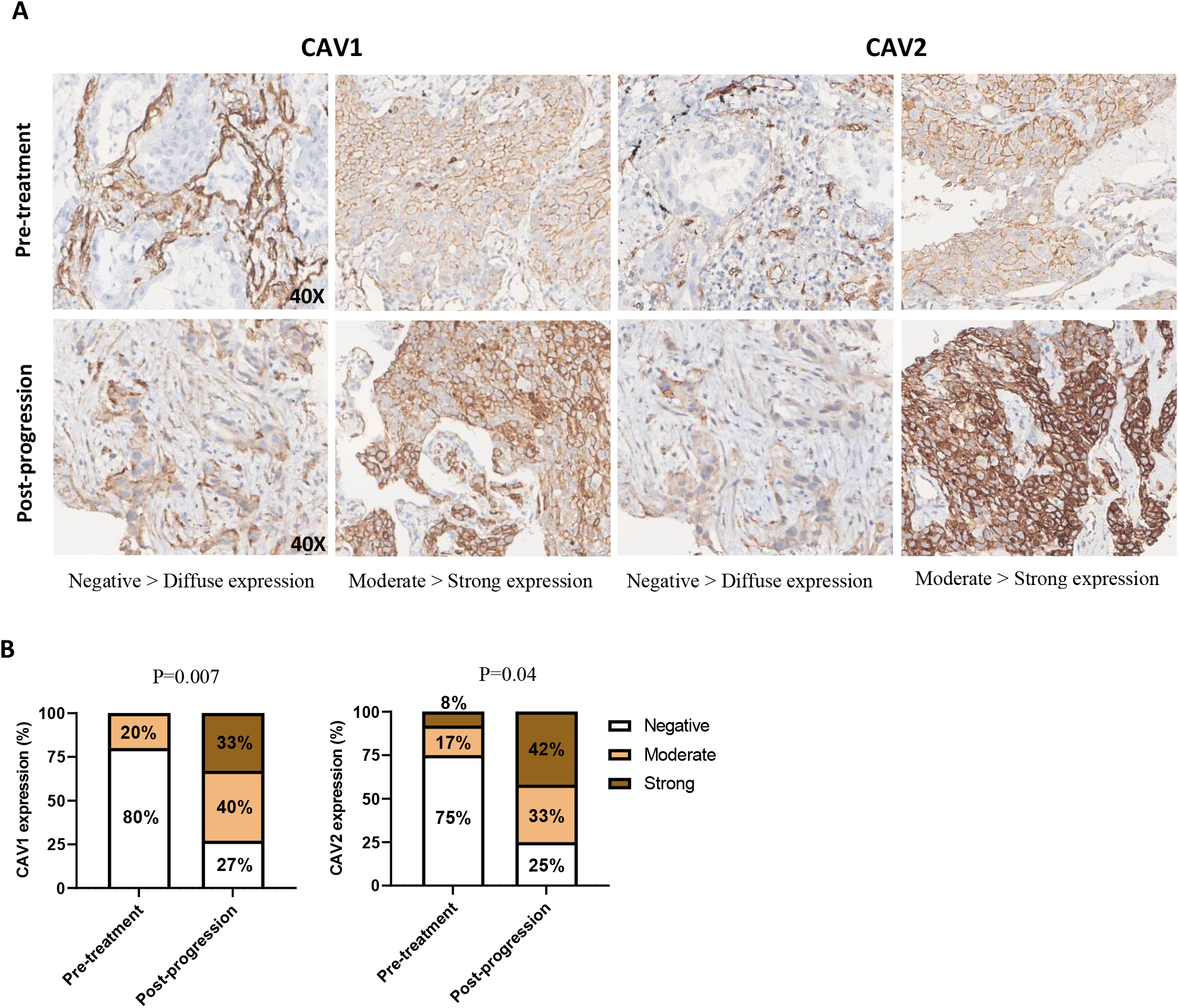
Increased expression of caveolins in EGFR-mutated human NSCLC after resistance acquisition to EGFR-TKI. (**A**) Microscopy images of CAV1 and CAV2 protein expression detected by immunohistochemistry in representative examples of untreated primary NSCLC (upper panel) and EGFR-TKI treated, p.T790M mutated relapsing tumors (lower panel) (magnification x400). CAV1 and CAV2 protein was mainly localized in the membrane with some cytoplasm staining in tumor cells. Endothelium and connective tissue also stained. (**B**) Percentage of pre-treatment and post-progression cases with negative, moderate, and strong expression of CAV1 and CAV2.

## DISCUSSION

One of the main implications of our findings is that mutation-driven resistance to targeted therapies can be transferred among cancer cells. Both *in vitro* and *in vivo* data show that in the presence of R cells, either using CM or through dual inoculation in the same animal, S cells acquire resistance faster. In mice, this results in shorter progression-free survival. Most importantly, we show that acquisition of resistance, as observed in S cells that grow alone (SI animals), is intrinsically different from what occurs in S cells that grow in DI animals. In the former, 70% of the tumours present with known EGFR-TKI resistance-associated mutations, namely MET and ERBB2 amplification, whereas in the latter these events are rare, with 90% of the tumours showing no evidence of mutation-associated resistance mechanisms.

Our results challenge the established paradigm that resistance to targeted therapies occurs because cancer cells harbouring resistance-conferring mutations, which initially exist in small numbers, are selected, accumulate, and eventually take over the entire tumour cell population through a vertical transmission model solely reliant on cancer cell division *(5)*. Furthermore, our clinical data on the AF of sensitizing and resistance mutations shows that in LC patients who relapse after EGFR-TKI treatment, the p.T790M mutation is present only in a fraction of the relapsing tumour cells. Assuming equal ploidy for the sensitizing and resistance alleles, as suggested by NGS CNV analysis, our data indicates that the fraction of cancer cells carrying the resistance mutation corresponds to about one-third, although this proportion may vary widely. These observations, support the idea that although resistance stems from a genetic mutation, cancer cells may become resistant even though they do not carry that mutation, opening the door to a model of cell-to-cell horizontal transfer of resistance.

Horizontal transfer of phenotypes is commonly observed in bacteria, which by exchanging genetic material, gain new traits such as resistance to antibiotics *(22)*. Horizontal transfer can also take place in eukaryotes, with exogenous DNA impacting the phenotype of recipient cells via mediators such as transposons, cfDNA, or extracellular vesicles (EVs) *(22)*. However, the role of horizontal transfer in cancer is largely unknown. Our results do not support the hypothesis that acquisition of resistance in S cells “exposed” to R cells is due to genetic transfer of the p.T790M mutation. Even though the resistance phenotype is stable in S cells, as these cells maintain the capacity to grow *in vitro* and *in vivo* under treatment with erlotinib, the p.T790M mutation originating in R cells could hardly be detected in S cells. Moreover, even if we accept the remote possibility that DNA coding for the p.T790M mutation is stably incorporated into the genome of some S cells, endowing them with resistance to erlotinib, this mechanism alone could not explain the widespread resistance phenotype observed in the resulting S tumours.

We reason that the full blown and stable resistance phenotype observed in our model is triggered by the p.T790M mutation in R cells, coupled with a continuous flow of molecular information from R to S cells, and among S cells. It has been shown that EVs carry proteins, RNA transcripts and other molecular material *(23, 24)*, making it a likely candidate to explain horizontal transfer of resistance across cancer cells. In fact, EV-mediated transfer of molecules has been shown to contribute to phenotypic reprogramming and functional re-education of recipient cells *(25, 26)*. Noteworthy, it has been recently reported that exosomal transfer of wild-type EGFR protein promotes resistance to osimertinib (a third-generation EGFR-TKI drug) in LC *(27)*.

A key finding of our study is the demonstration that horizontal transfer of resistance triggers a change in the transcriptional landscape of ERS tumours. Pathway analysis showed significant changes in genes involved in endocytosis, namely overexpression of CAV1 and CAV2. Overexpression was confirmed at the mRNA and protein level in samples from our *in vivo* model and at the protein level in clinical LC samples. The results obtained in clinical samples are especially relevant because they correspond exactly to the model of acquisition of resistance to erlotinib determined by the occurrence of the p.T790M EGFR mutation. In our series, 67% and 60% of the cases showed gain of CAV1 and CAV2 expression, respectively, comparing the pre-treatment to the post-progression samples.

Caveolins are membrane-associated proteins that play a key role in the formation of non-planar lipid rafts known as caveolae *(28)*. In turn, caveolae are important for signal transduction through their capacity to selectively concentrate proteins, such as membrane receptors, kinases, and phosphatases, thereby promoting specific molecular interactions *(29)*. Caveolins are also involved in intracellular trafficking, including endocytosis, through their localization in distinct cellular organelles and compartments *(30)*. Caveolins, namely CAV1 and CAV2, have been involved in cancer with both oncogenic and tumour suppressor properties being described *(31)*. CAV1, for instance, tends to be expressed in normal cells, downregulated during neoplastic transformation and first stages of tumour progression, and re-expressed in late stages associated with treatment resistance and metastasis *(31)*.

In LC, increased CAV1 expression was associated with gemcitabine resistance *(32)*, and poor prognosis *(33, 34)*. Caveolins have also been demonstrated to modulate EGFR activity in cancer cells. CAV2 was shown to promote the growth of renal cell carcinoma cells through EGFR signalling *(35). In vitro*, CAV1 was shown to increase LC cell proliferation, migration, and invasion through EGFR phosphorylation *(36)*. Most importantly, silencing of CAV1 in LC cells led to enhanced sensitivity to EGFR-TKI drugs by down-regulating phosphorylation of EGFR *(37)*, directly supporting our contention that the acquisition of resistance to erlotinib is associated with increased CAV1 and CAV2 expression. The precise mechanism linking caveolins to EGFR-TKI resistance is not known, however, published evidence indicates it is likely to involve increased EGFR signalling and increased endocytosis. There is evidence that such increase in endocytosis is accompanied by overexpression and nuclear translocation of both EGFR and other membrane receptors *(30, 38)*, which in turn may provide survival signals to overcome the inhibitory effect of EGFR-TKI drugs.

In conclusion, our results suggest that the EGFR-TKI resistance phenotype can be transferred from resistant to sensitive cells both *in vivo* and *in vitro*, resulting in reprogramming of the cells that acquire resistance. Our findings point out a key role for endocytosis, coupled with caveolae-associated regulation of signaling and intracellular trafficking, in the acquisition of resistance to targeted therapies in cancer. Even though our study does not elucidate how transfer of resistance occurs, or the specific mechanism linking caveolins with EGFR-TKI resistance, we present a strong case to challenge the current paradigm of cell division and selection as the sole mechanism underlying transmission of mutation-driven resistance to targeted therapies. In addition, our study provides a proof of concept to trigger further research on whether these results can be applied to other models of targeted therapy and mutation-driven acquisition of resistance in cancer.

## MATERIALS AND METHODS

### Human samples

Samples from patients with advanced lung adenocarcinoma (unresectable stages IIIB and IV) were obtained from the Pulmonology Departments of Centro Hospitalar de São João and Centro Hospitalar de Vila Nova de Gaia/Espinho. Pre- and post-EGFR-TKI treatment tumour tissue biopsies from 15 LC patients were used for immunohistochemistry. Post-EGFR-TKI treatment liquid and tissue biopsies from LC patients (14+14) were selected for NGS analysis. Blood samples were collected in K_2_EDTA tubes (BD Vacutainer® PPT™ Plasma Preparation Tube, Becton Dickinson, Franklin Lakes, USA). The plasma fraction was separated from the blood cells by centrifugation at 1200xg during 10min. The collected plasma was aliquoted and stored at −80°C for cell-free DNA (cfDNA) extraction. Histology and cytology specimens were formalin-fixed and paraffin-embedded and reviewed by a pathologist.

### Cell lines and cell culture

Human LC cell lines HCC827 (ATCC Cat# CRL-2868™) and H1975 (ATCC Cat# CRL-5908™) were cultured in RPMI-1640 medium [GIBCO, USA], supplemented with 10% (v/v) foetal bovine serum (FBS) [GIBCO, USA], 100 U/mL penicillin and 100μg/mL Streptomycin [GIBCO, USA]. Cells were kept at 5% CO_2_ and 37°C in a humidified atmosphere. Cells were tested for mycoplasma during our study and STR profiled.

### Conditioned medium experiment

LC cells sensitive to the EGFR-TKI (S cells) were seeded in a 6-well plate, 2.5×10^5^ cells/well. After 24h, medium from LC cells resistant to the EGFR-TKI (R cells) was collected, centrifuged at 4,000rpm for 10min and filtered using 0.45μm filter, before being added to S cells, together with 0.01μM EGFR-TKI. The EGFR-TKI included in this study was erlotinib [LC Laboratories, Woburn, MA, USA]. Erlotinib stock solution was prepared in 0.5% [wt/vol] methylcellulose [Sigma-Aldrich®, St. Louis, MO, USA] and 0.4% [vol/vol] Tween-80 [Sigma-Aldrich®, St. Louis, MO, USA] solution. Experiments were performed with three replicates per condition.

### LC xenograft model

S cells (5 × 10^6^) and R cells (1× 10^5^) were subcutaneously injected into the dorsal flank of 8-12-weeks-old immunodeficient C57BL/6 *Rag2*^*−/−*^*;Il2rg*^*−/−*^ mice. We engrafted S cells in one flank and once tumours reached a size ≈800mm^3^, R cells were engrafted in the opposite flank. When the S tumours reached ≈1000mm^3^, erlotinib was administered (3 times/week, oral gavage, 25mg/kg). Due to rapid growth of the R tumours, surgery was performed to remove these tumours when it reached 1500mm^3^ of size and after 1-2 weeks of recovery, mice were again engrafted with R cells. This procedure allowed to follow-up S cell tumours for longer periods of time until acquisition of resistance to erlotinib. In both groups, animals were sacrificed at humane endpoint. A set of animals were engrafted with S cells in a single flank and used as control. Tumour size was measured with callipers every other day and tumour volume calculated by the formula length×width^2^/2. <u>Disease</u> progression was defined as an increase of at least 75% in volume with respect to the minimum tumour volume after treatment. At humane endpoint, mice were euthanized, and tumours dissected and immersed in 10% formalin, for histological and molecular analysis.

### DNA extraction

DNA from cells was extracted using DNeasy® Blood & Tissue kit [QIAGEN, Hilden, Germany], according to manufacturer’s instructions. Tumour samples from mice and tissue biopsies from patients were cut in 10μm sections using the Microm HM 335 E paraffin microtome [GMI, Ramsey, MN, USA] and 3 sections of each sample were collected. DNA and RNA extraction were performed using MagMAX™ FFPE DNA/RNA Ultra Kit [Thermo Fisher Scientific, Waltham, MA, USA], according to manufacturer’s instructions.

cfDNA from plasma patients was extracted using MagMAX Cell-Free DNA Isolation kit [Applied Biosystems, Life technologies, Waltham, MA, USA], according to manufacturer’s instructions. Fragment distribution and concentration of DNA from plasma was evaluated using 2200 TapeStation [Agilent Technologies, Santa Clara, CA, USA], according to manufacturer’s instructions. The results were analysed with the TapeStation Analysis Software [Invitrogen, Life technologies, Waltham, MA, USA]. All plasma DNA quantifications were normalized for volume of plasma collected [cfDNA concentration (ng/mL of plasma) = cfDNA quantification (ng/μL) x elution volume (μL) / plasma volume (mL)].

DNA and RNA concentration were determined using Qubit® 2.0 Fluorometer [Invitrogen, Life technologies, Waltham, MA, USA], double stranded DNA high sensitivity (HS) or RNA HS assay, according to manufacturer’s instructions.

### DNA fingerprinting

DNA from cells was amplified through PCR with primers for the following locus: Penta E, D18S51, D21S11, THO1, D3S1358, FGA, TPOX, D81179, vWA, Amelogenin, Penta D, CSF1PO, D16S539, D7S820, D14s317 and D5S818 with the Powerplex 16 HS system, that allows the co-amplification and simultaneous detection of the 16 described loci. The amplified fragments were then detected with capillary electrophoresis using the 3500 Genetic Analyser sequencer [Applied Biosystems] and the genotypes were assigned with the GeneMapper v5.0 [Applied Biosystems].

### Next Generation Sequencing (NGS)

NGS libraries were prepared using the Oncomine™ Lung cfDNA Assay for mouse xenografts and for human liquid biopsies, and the Oncomine™ Focus Assay for human tissue biopsies according to the manufacturer’s instructions [Thermo Fisher Scientific, Waltham, MA, USA]. The resulting libraries were purified using Agencourt AMPure XP [Beckman Coulter] and quantified by qPCR using the Ion Library TaqMan® Quantitation Kit [Thermo Fisher Scientific, Waltham, MA, USA], according with the manufacturer instructions. The quantified stock libraries were then diluted to 50pM, pooled and loaded onto Ion 530™ or 540™ chips using Ion Chef™ for templating [Thermo Fisher Scientific, Waltham, MA, USA] and the loaded chips were then sequenced in a S5XL sequencer. The sequencing quality was assessed through the coverage analysis plugin and the samples were analysed with Ion Reporter 5.6. Raw data was processed automatically on the Torrent Server™ and aligned to the reference hg19 genome.

### Digital PCR

TaqMan Mutation Detection Assays [Thermo Fisher Scientific, Waltham, MA, USA] were used on a Quantstudio 3D digital PCR system [Thermo Fisher Scientific, Waltham, MA, USA] to confirm variants with low allelic frequencies. This approach was used to detect specific alterations, with specific probes for each mutation, namely the Hs000000029_rm (for *EGFR* p.T790M mutation), <u>Hs000000026_rm</u> (for *EGFR* p.L858R mutation), <u>Hs000000027_rm</u> (for *EGFR* p.E746-A750 deletion) and <u>Hs00001004_mu</u> (for TP53 p.R273H mutation). The results were analysed with QuantStudio™ 3D AnalysisSuite™ Software.

### RNA-Seq

The integrity and quality of RNA was analysed using the Agilent 2100 Bioanalyzer [Agilent Technologies, Santa Clara, CA, USA]. RNA was converted to cDNA using the SuperScript VILO cDNA Synthesis Kit [Thermo Fisher Scientific, Waltham, MA, USA]. cDNA was amplified and library constructed with Ion AmpliSeq Transcriptome Human Gene Expression Kit [ThermoFisher Scientific, Waltham, MA, USA]. Samples were then prepared for deep sequencing using the Ion Chef™ System [Thermo Fisher Scientific, Waltham, MA, USA], loaded into the Ion 550 Chip, and sequenced using the Ion S5 XL Sequencer [Thermo Fisher Scientific, Waltham, MA, USA]. Sequencing quality was assessed through the plug-in coverage analysis and samples were analysed on the torrent Suite™ Software [Thermo Fisher Scientific, Waltham, MA, USA]. Differentially expressed genes between groups were identified using the Transcriptome Analysis Console Software v4.0.2 [Thermo Fisher Scientific, Waltham, MA, USA] using a double threshold based on fold-change (>=1 ou <=-1) and statistical significance of the change with FDR p-value <=0.05. DAVID webtool was used to determine significantly enriched gene ontology terms and pathways.

### Real time PCR

RNA was converted to cDNA using the SuperScript VILO cDNA Synthesis Kit [Thermo Fisher Scientific, Waltham, MA, USA], according to manufacturer’s instructions. The reactions were carried out in a StepOnePlusTM qPCR Real-Time PCR machine, in a volume of 10μL containing 1x TaqManTM Fast Advanced Master mix [Applied Biosystems], with 1x TaqManTM Advanced Assays probes Hs00971716_m1 and Hs00184597_m1 [Applied Biosystems] specific for the *CAV1* and *CAV2* gene, respectively, and 2.5μL cDNA. For mRNA expression normalization, two housekeeping controls were used: HPRT1 and GUSB [Applied Biosystems]. The 2^-ΔΔCT^ method was applied to analyse the relative change in gene expression. Three technical replicates were made for each sample.

### Immunohistochemistry

Tissue sections were obtained in coated slides [Thermo Scientific, Waltham, MA, USA], 3μm each section, using the Microm HM 335 E paraffin microtome [GMI, Ramsey, MN, USA] followed by incubation at 65°C for 1h. Each section was probed overnight at 4°C with an optimized concentration of the anti-CAV1 (Sigma-Aldrich®, catalog #HPA049326, 1:1000) and anti-CAV2 (Novus Biologicals, catalog #NBP2-98731, 1:500) polyclonal antibodies diluted in antibody diluent [Thermo Scientific, Waltham, MA, USA]. CAV1 and CAV2 proteins were detected by peroxidase-DAB (diaminobenzidine) chemistry using the REAL EnVision detection system kit [Dako, Glostrup, Denmark]. CAV1 and CAV2 immunoreactivity was classified as: (1) negative, when tumour cells showed complete loss of expression; (2) moderate, when less than 50% of the tumour cells showed preserved membrane expression, or, irrespective of the percentage of cells, weaker membrane staining compared with the control; or (3) strong, when more than 50% of the tumour cells showed preserved membrane expression.

### Statistical analysis

All statistical analyses were performed with GraphPad Prism [GraphPad Prism, RRID:SCR_002798], SPSS Statistics V27 Release 27.0.1.0. [IBM, RRID:SCR_019096] or R software package (https://www.r-project.org). The data were showed as means ± standard deviation (SD) and analysed by Student’s *t*-test analysis unless otherwise indicated. Log-Rank (Mantel-Cox) test was used to compare progression-free survival onset between different mice models. Pearson correlations between gene expressions of CAV1 and CAV2 were performed using the “cor.test” package in R.

## List of Supplementary Materials

Figs. S1 to S3.

Tables S1 and S2.

## Acknowledgments

We thank Dr Nuno Alves, i3S, Porto, Portugal and Dr James Di Santo, Institute Pasteur, Paris, France, for kindly gifting the Ilrg2-/-. We thank Dr. Dina Leitão for assistance in the immunohistochemistry staining. Elaboration of figures 1A and 2A was performed using BioRender.com.

## Author contributions

Conceptualization: SJN, SAM, JLC and JCM

*In vitro* and *in vivo* experiments: SJN, ARO, JFM, MS, BA

Other methodology: SJN, ARO, JFM, JP, CSM, JCM, PO.

Analysis of results: SJN, ARO, JFM, PO, LP, BC, JP, CSM, JLC, JCM

Selection of clinical samples and analysis of clinical data: AB, VH, MGOF, CSM, JLC, JCM

Funding acquisition: JLC, JCM.

Writing - original draft: SJN, JCM.

Writing – review & editing: all authors

## Data and materials availability

All data, except for RNASeq raw data, are available in the main text or the supplementary materials. The RNASeq datasets used and/or analyzed during the current study are available from the corresponding author upon request.

## Ethics approval and informed consent

The study was conducted according to the guidelines of the Declaration of Helsinki. Ethical review and approval were waived as the study is in accordance with Article 19 (“DNA Banks and Other Biological Products”) of Portuguese Law No. 12/2005 of 26 January (“Personal genetic information and health information”), which states that in the case of using retrospective samples from human origin or in special situations where the consent of the people involved cannot be obtained because of the amount of data or subjects, of their age, or another reason comparable, material and data can be processed but only for purposes of scientific research or epidemiological and statistical data collection.

Animal studies were approved by the national authority Direção Geral de Alimentação e Veterinária (DGAV reference 0421/000/000/2017) and reviewed by the i3S Animal Welfare and Ethics Body (reference 2016/22). All mice were housed under standard housing conditions at the i3S animal facility.

