## Supplemental tables and figures for "Resistance to EGFR inhibitors in lung cancer occurs through horizontal transfer and is associated with increased caveolins expression"

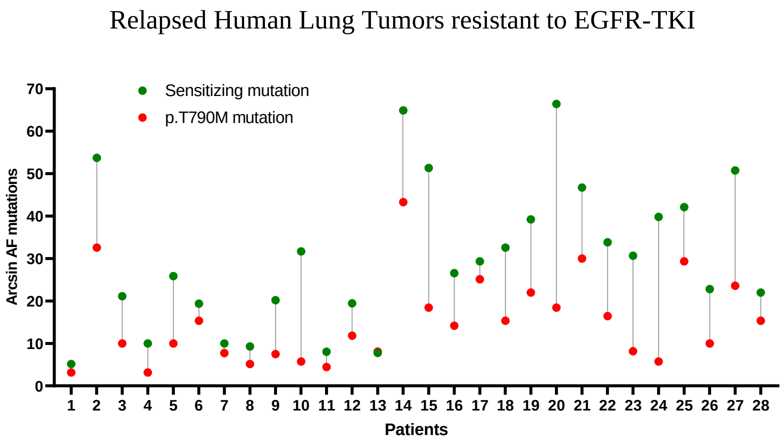


**Figure S1.** Allele frequencies determined by NGS in samples from lung cancer patients after disease progression. Arcsin square root transformed percentages of allele frequency of sensitizing mutation (green dots) and p.T790M resistance mutation (red dots).

**S tumor** | **R tumor**

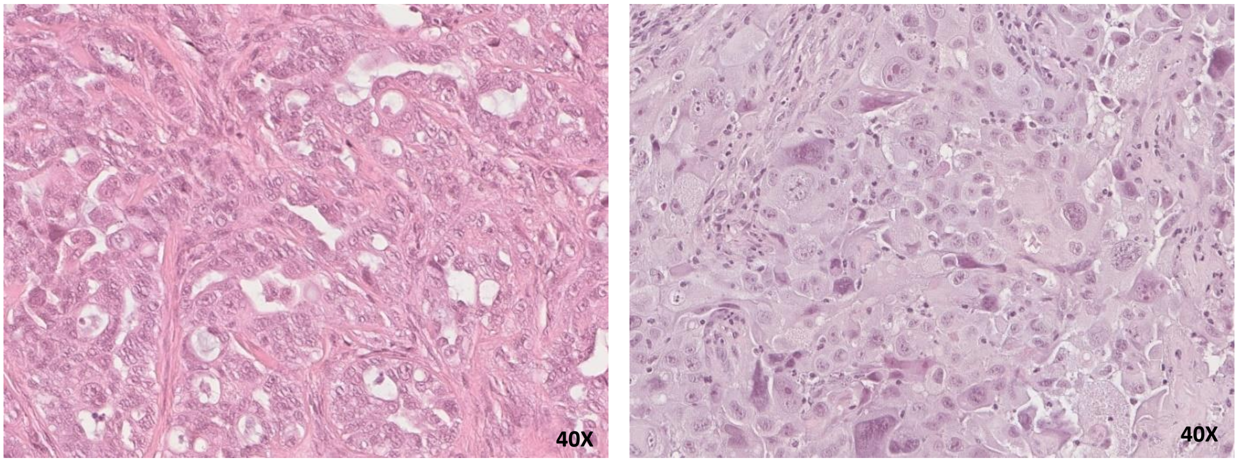


**Figure S2.** Histology of S and R tumors from DI mice. Left image (S tumor) - malignant neoplasia with characteristics of carcinoma, consisting of large cells and acinar/cribriform components. Right image (R tumor) - malignant neoplasia with characteristics of solid pattern carcinoma, consisting of large polygonal cells with eosinophilic cytoplasm and accentuated nuclear pleomorphism (magnification x400).

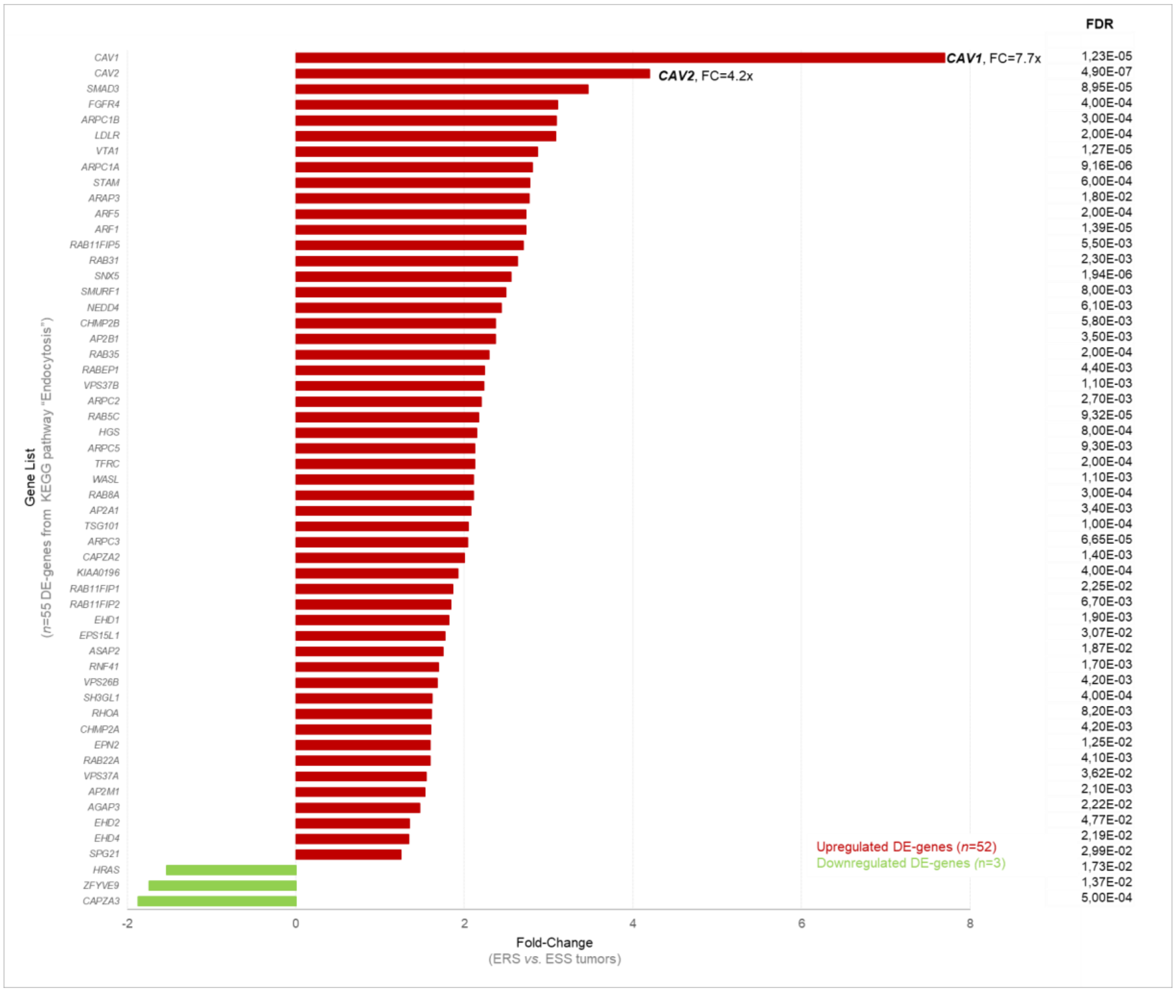


**Figure S3.** Differentially expressed (DE) genes between ERS and ESS tumors from endocytosis KEGG pathway. 55 DE-genes from endocytosis pathway of which 52 are upregulated and 3 downregulated (FDR p-value < 0.05).

**Table S1.** Detection of sensitizing (p.E746_A750del) and resistance (p.T790M) *EGFR* mutations in S tumors from DI animals and S tumors from SI animals, detected by NGS.

**DI ANIMALS**

| SAMPLE ID | EGFR mutation | AF (%) |
|---|---|---|
| 1 | p.E746_A750del<br>p.T790M | 95,5<br>0,1 |
| 2 | p.E746_A750del<br>p.T790M | 90,9<br>0,1 |
| 3 | p.E746_A750del<br>p.T790M | 90,0<br>0,1 |
| 4 | p.E746_A750del | 93,4 |
| 5 | p.E746_A750del | 96,8 |
| 6 | p.E746_A750del | 86,2 |
| 7 | p.E746_A750del | 94,9 |
| 8 | p.E746_A750del | 91,3 |
| 9 | p.E746_A750del<br>p.T790M | 93,7<br>0,1 |
| 10 | p.E746_A750del<br>p.T790M | 91,9<br>0,1 |
| 11 | p.E746_A750del | 86,4 |
| 12 | p.E746_A750del | 96,7 |
| 13 | p.E746_A750del | 93,9 |
| 14 | p.E746_A750del | 94,3 |
| 15 | p.E746_A750del | 96,6 |
| 16 | p.E746_A750del | 93,3 |
| 17 | p.E746_A750del | 93,6 |
| 18 | p.E746_A750del | 95,0 |
| 19 | p.E746_A750del | 93,7 |
| 20 | p.E746_A750del<br>p.T790M | 93,9<br>0,1 |

**SI ANIMALS**

| SAMPLE ID | EGFR mutation | AF (%) |
|---|---|---|
| 1 | p.E746_A750del | 86,1 |
| 2 | p.E746_A750del | 90,4 |
| 3 | p.E746_A750del | 91,2 |
| 4 | p.E746_A750del | 91,1 |
| 5 | p.E746_A750del | 93,0 |
| 6 | p.E746_A750del | 91,5 |
| 7 | p.E746_A750del | 90,0 |
| 8 | p.E746_A750del | 97,2 |
| 9 | p.E746_A750del | 94,7 |
| 10 | p.E746_A750del | 92,7 |
| 11 | p.E746_A750del | 94,9 |
| 12 | p.E746_A750del | 94,1 |

**Table S2.** CAV1 and CAV2 expression in pre-treatment and post-progression samples.

| Case | CAV1 pre-progression | CAV1 post-progression | CAV2 pre-progression | CAV2 post-progression |
|---|---|---|---|---|
| 1 | Negative | Strong | Negative | Strong |
| 2 | Negative | Strong | Negative | n.a. |
| 3 | Negative | Negative | Negative | n.a. |
| 4 | Negative | Moderate | Negative | Moderate |
| 5 | Negative | Negative | n.a. | n.a. |
| 6 | Negative | Strong | n.a. | Strong |
| 7 | Negative | Moderate | Negative | Negative |
| 8 | Moderate | Strong | Strong | Strong |
| 9 | Negative | Negative | Negative | Negative |
| 10 | Moderate | Strong | Moderate | Strong |
| 11 | Negative | Moderate | n.a. | Moderate |
| 12 | Negative | Moderate | Negative | Moderate |
| 13 | Negative | Moderate | Negative | Moderate |
| 14 | Moderate | Moderate | Moderate | Strong |
| 15 | Negative | Negative | Negative | Negative |

n.a. not available (no tumor representation)
